# *Citrus* IntegroPectin bioconjugates induce apoptosis, decrease proliferation, reduce reactive oxygen species, alleviate mitochondrial oxidative stress, upregulate miR-146 expression and downregulate expression of Interleukin-8 in lung cancer cells

**DOI:** 10.1101/2025.10.03.680243

**Authors:** Claudia D’Anna, Caterina Di Sano, Giovanna Li Petri, Giuseppe Angellotti, Simona Taverna, Giuseppe Cammarata, Rosaria Ciriminna, Mario Pagliaro

## Abstract

Tested on adenocarcinoma A549 cell line, IntegroPectin bioconjugates derived from the industrial processing waste of different organically grown *Citrus* fruits exert a pleiotropic activity in their anticancer mechanism by targeting different molecular pathways. This study shows that these novel bioconjugates induce apoptosis, decrease short-term proliferation response, reduce reactive oxygen species, alleviate mitochondrial oxidative stress, upregulate miR-146 expression and downregulate expression of Interleukin-8. Given the health beneficial properties of both citrus pectin and citrus flavonoids, and the frequency of lung cancer, said findings support urgent investigation of these new flavonoid-pectin conjugates in preclinical and clinical cancer treatment trials.

## 1. Introduction

Lung cancer is the main cause of cancer-related mortality,^[1]^ with non-small-cell lung cancer (NSCLC) being the most common form (85% share) of lung tumor. Owing to late-stage detection, life expectancy is poor. Median overall survival was only 1 year after diagnosis in 2015 when 1.6 million of deaths occurred worldwide.^[2]^ Research aimed to develop new therapies for NSCLC ranges from adoptive cell transfer and its forms including tumor-infiltrating lymphocytes therapy,^[3]^ through new drugs enhancing the immune system’s ability to fight cancer by blocking the proteins cancer cells use to evade immune cells,^[4]^ to natural products able to modulate the immune system, drive apoptosis, and arrest cancer cell proliferation^[5]^ and prevent metastatic spread.^[6]^

In this context, we recently reported that *Citrus* IntegroPectin bioconjugates derived via acoustic cavitation (AC) from the processing waste of lemon, red orange, and sweet orange, inhibit cancer cell migration and long-term proliferation in the adenocarcinoma A549 cell line.^[7]^ Both long-term proliferation and cell migration are crucial processes in cancer progression,^[8]^ with epithelial-mesenchymal transition (EMT) being a process by which epithelial cells acquire mesenchymal traits, enhancing.

Sourced via AC or via hydrodynamic cavitation (HC) in water only of the fresh residue of industrial manufacturing of *Citrus* juice sourced from different organically grown fruits, IntegroPectin is a family of bioconjugates of low-methoxyl (LM) pectin and flavonoids.^[9]^

Sourced from orange, lemon, grapefruit and other *Citrus* fruit processing waste, these bioproducts exert broad and pronounced bioactivity including antioxidant, anti-inflammatory, cardioprotective, neuroprotective, mitoprotective, antimicrobial and anticancer properties demonstrated through both *in vitro* and *in vivo* experiments.^[9]^

The vastly enhanced biological activity of *Citrus* IntegroPectin in comparison to commercial lemon (and orange) pectin has been ascribed to its unique molecular structure (ultralow degree of methylation, abundant rhamnogalacturonan-I (RG-I) regions and complete decrystallization of the homogalacturonan (HG) regions due to cavitation destroying the “fringed-micellar” structure of the crystalline regions of the semicrystalline pectin biopolymer, coupled to the abundance of highly bioactive flavonoids (and terpenes) at its surface.^[9]^

In this study, we investigate the anticancer properties of *Citrus* IntegroPectin bioconjugates derived from the processing waste of lemon, red orange, and sweet orange, by assessing their ability to induce apoptotic cell death, decrease short-term proliferation response, reduce reactive oxygen species (ROS) and alleviate mitochondrial stress in adenocarcinoma lung cancer cells A549.

One key feature of cancer cells is their ability to evade apoptosis, which allows them to survive despite DNA damage, oxidative stress, and other cytotoxic insults. In detail, cancer cells avoid apoptosis in many ways, including inhibition of caspases and overexpression of anti-apoptotic B cell lymphoma-2 (BCL2) family of proteins observed in abut 50% of malignancies, independent of subtype.^[10]^

Another molecular pathway representing a promising avenue for cancer therapy is overexpression of miR-146a (a microRNA) that in lung cancer inhibits cell growth, cell migration and induces apoptosis in NSCLC cells.^[11]^ MicroRNAs (miRNAs) are a class of small non-coding ribonucleic acids that regulate gene expression in later stages of the translation process by binding to genomic regulatory sites.

In cancer, uncontrolled cell migration is due to the elevated ROS level, which promotes DNA mutations, uncontrolled cell proliferation, and resistance to cell death.^[12]^ Oxidative stress occurs when there is an imbalance between the production of ROS and the antioxidant defense capacity of the cell, leading to excess ROS. ROS play an essential physiological role in many cellular processes, such as in cellular signaling and immune response (“oxidative eustress”.^[13]^ In brief, moderate ROS levels support cancer cell survival, whereas high ROS levels can induce apoptosis, thereby establishing a dual role that today makes ROS a target for cancer therapy.^[14]^

In the case of lung cancer, ROS promote cancer origination through oxidative stress and base-pair substitution mutations in pro-oncogenes and tumor suppressor genes, at later stages they help the cancer cells in invasion and metastasis by activating the nuclear factor kappa beta (NF-κB, enhancing transcription of anti-apoptotic genes) and mitogen-activated protein kinase (MAPK) signaling pathways, whereas at advanced stages, they promote cell cycle arrest inducing apoptosis in cancer cells.^[15]^

## 2. Results

Photographs in Fig.S1 show the freeze-dried *Citrus* IntegroPectin samples sourced via AC from industrial biowaste of lemon, sweet orange, and red orange obtained from citrus juice production used throughout this work. The IntegroPectin bioconjugates used in this work underwent further purification via membrane dialysis to remove residual sugars. We briefly remind that these flavonoid-pectin bioconjugates consist of highly de-esterified *Citrus* pectins rich chiefly in citrus flavonoids such as hesperidin, naringin, eriocitrin and kaempferol, containing also minor amounts of phenolic acids such as *p*-coumaric acid.^[8]^

Photographs in Fig.S2 show lemon IntegroPectin before and after 24 h of dialysis. The effect of dialysis on the characteristics of each IntegroPectin sample was clearly beneficial. The dialyzed IntegroPectin bioconjugates show no hygroscopic behavior even after one week of storage at 4 °C. Furthermore, all dialyzed IntegroPectin samples exhibit a noticeably brighter color compared to the untreated materials, indicating further concentration of flavonoids and other bioactive compounds. Indeed, the amount of hesperidin (Table S1) was exceptionally high in red orange (62.31 mg/g) and also in sweet orange (40.35 mg/g) IntegroPectin bioconjugates, namely nearly one order of magnitude higher when compared to the non-dialyzed analogues (7.2 mg/g in red orange and 6.8 mg/g for sweet orange IntegroPectin conjugates).^[8]^

Displaying the characteristic IR signals of *Citrus* IntegroPectin bioconjugates sourced via HC from lemon, red orange, bitter or sweet orange,^[16]^ the FTIR spectra (Fig.1) are very similar to those of the non-dialyzed IntegroPectin samples, indicating highly de-esterified pectins rich in flavonoids and phenolic acids.^[7]^ The strongf bands in the 1550–1800 cm^−1^ region, with maxima at 1613 and 1720 cm^-1^, are assigned to the stretching modes of carboxylate groups (ν_as_COO^−^) and esterified galacturonic acid (νC=O_ester_), respectively. In this region, the relative intensities of the ester-related components are much higher for lemon and sweet orange suggesting a higher degree of esterification than red orange IntegroPectin.

**Figure 1.**
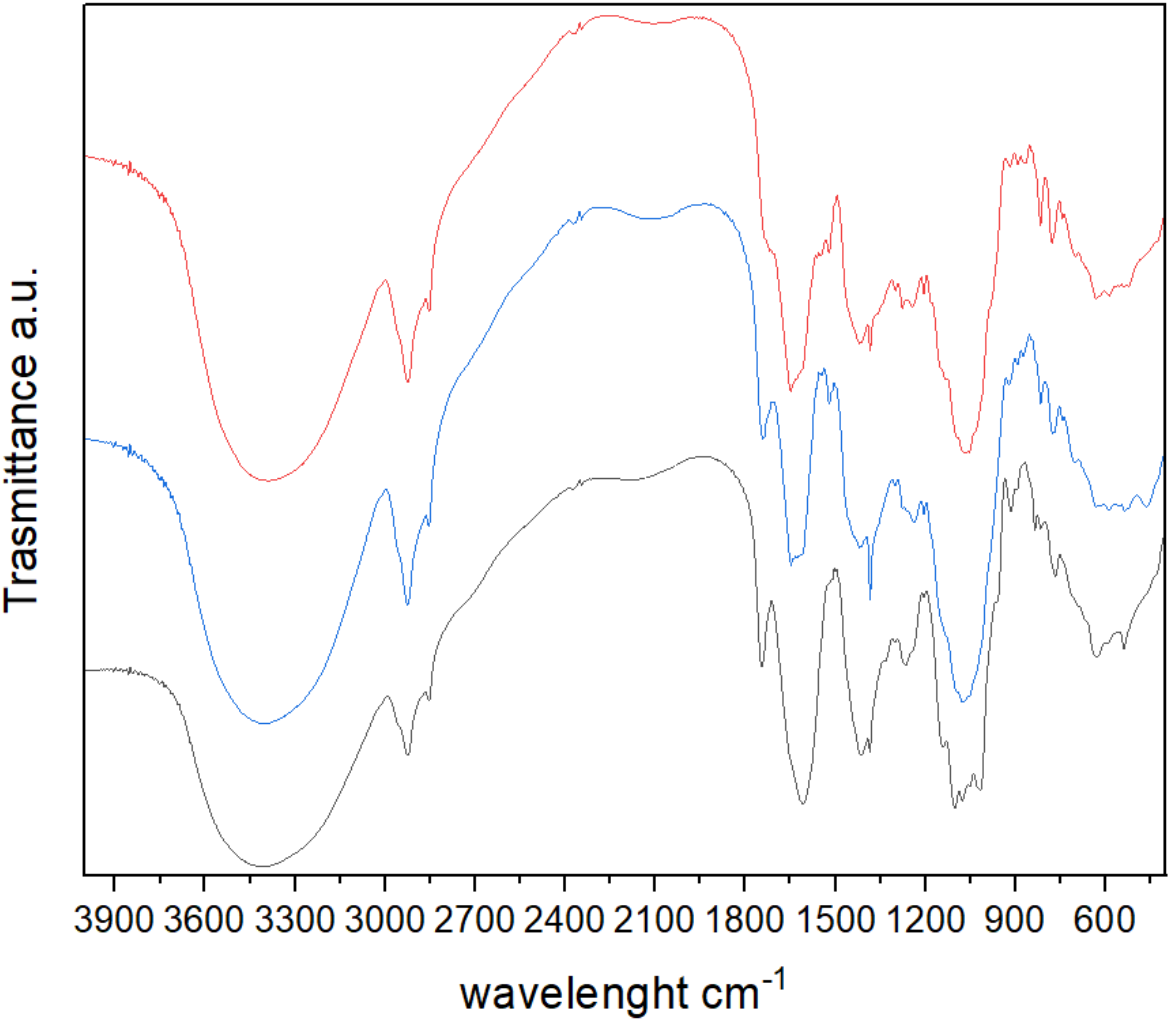
FTIR spectra of dialyzed *Citrus* IntegroPectin samples extracted from lemon (black), sweet orange (blue) and red orange (red).

All samples exhibit in the 950–1200 cm^−1^ region contains very intense bands typical of pectin assigned to skeletal stretching modes (νC-C and νC-O-C) of the pyranose rings and of the glycosidic bonds, and to a combination of the νC-OH and νC-C modes from the pyranose rings, indicating preservation of the pectin structure in all three extracts. The signal at 2931 cm^-1^ is due to the stretching vibrations of C-H bonds in CH and CH_2_ of pyranose rings in the polysaccharide chain, whereas the broad band at 3406 cm^-1^ is due to the νO–H band stretching vibration of the alcohol groups in the pyranose ring and of adsorbed water. The signal is similar in all three IntegroPectin sourced via AC whereas the lack of high wavenumber shoulder (at ∼3500 cm^−1^) in all samples shows evidence of similar lack of OH groups with weaker hydrogen-bond interactions.^[17]^

### 2.1 Effects on cell cycle in A549 cell line

A549 cells were cultured for 24 h with IntegroPectin bioconjugates dissolved in PBS at 0.5 and 1.0 mg/mL load. Cell cycle was assessed by cytofluorimetric analysis. As shown in Fig.3 all three IntegroPectin phytocomplexes promote apoptosis at all tested concentrations. Specifically, lemon IntegroPectin at a concentration of 1.0 mg/mL and red orange IntegroPectin at concentrations of 0.5 and 1 mg/mL induce a significant increase in apoptotic cell death.

**Figure 3.**
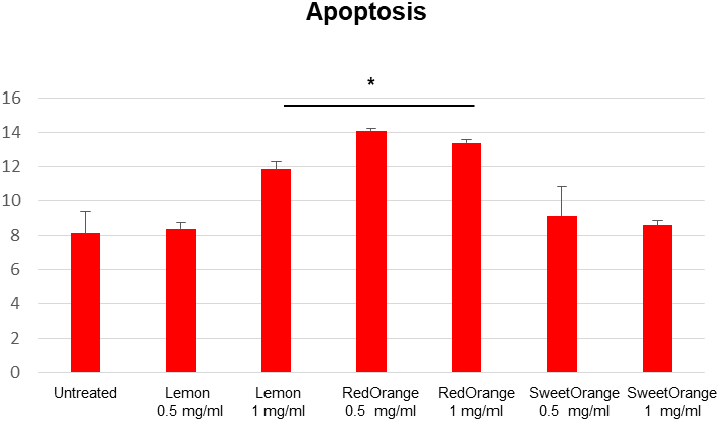
Apoptosis of A549 cells treated with lemon, red orange, sweet orange IntegroPectin at different concentrations.

The cell cycle analysis (plots in Fig.4) reveals that this apoptotic response is accompanied by an arrest at the G0/G1 phase. This arrest in the G0/G1 phase is a crucial event, as it effectively halts the progression of tumor cells into the S and G2/M cell proliferation phases, where DNA replication and mitosis occur.

**Figure 4.**
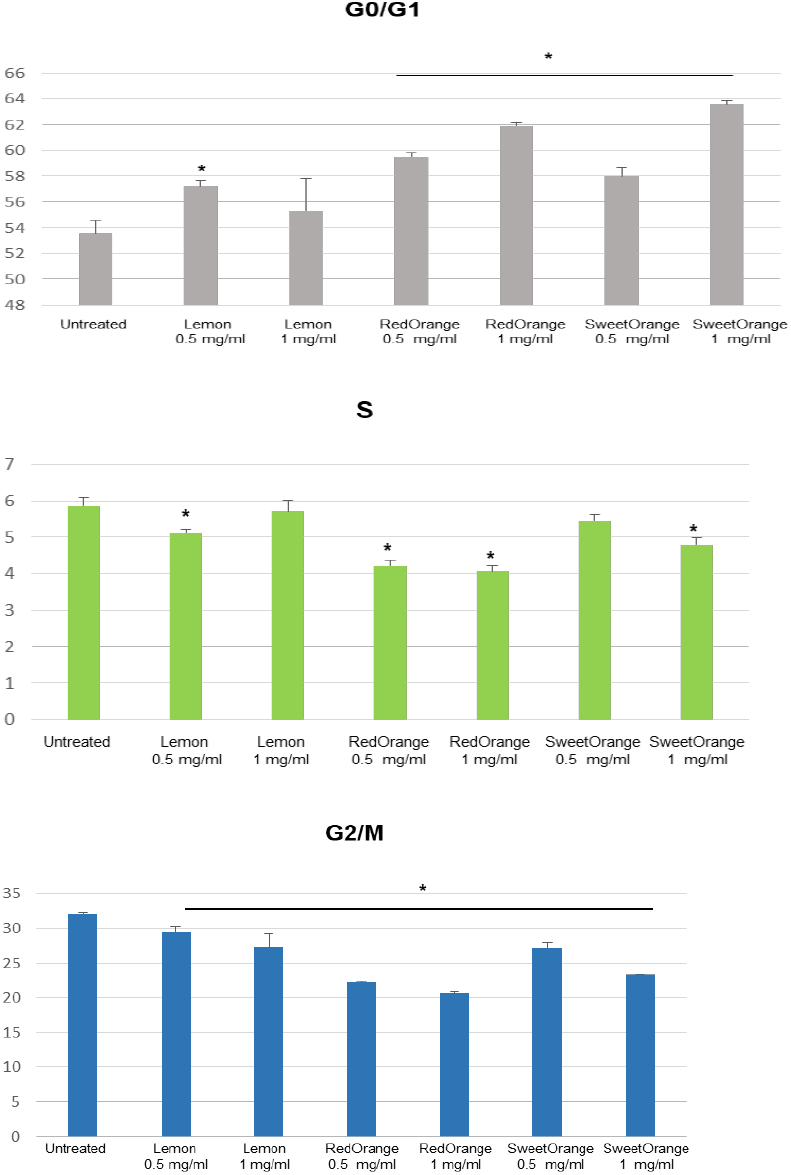
Cell cycle analysis for A549 cells treated with lemon, red orange, sweet orange IntegroPectin at different concentrations.

### 2.2 Effects on A549 cell line short-term proliferation

Following treatment with different IntegroPectin bioconjugates, short-term proliferation was assessed by cytometry analysis flow using carboxyfluorescein diacetate succinimidyl ester (CFSE) cell-permeable dye. Higher fluorescence intensity indicates fewer cell divisions.

As shown in Fig.5, at all tested concentrations the flow cytometry analysis revealed a significantly increased CFSE signal, consistent with reduced proliferation (as cells divide, the CFSE is progressively diluted, leading to a decrease in fluorescence intensity. A higher initial fluorescence intensity suggests a lower number of cell divisions).

**Figure 5.**
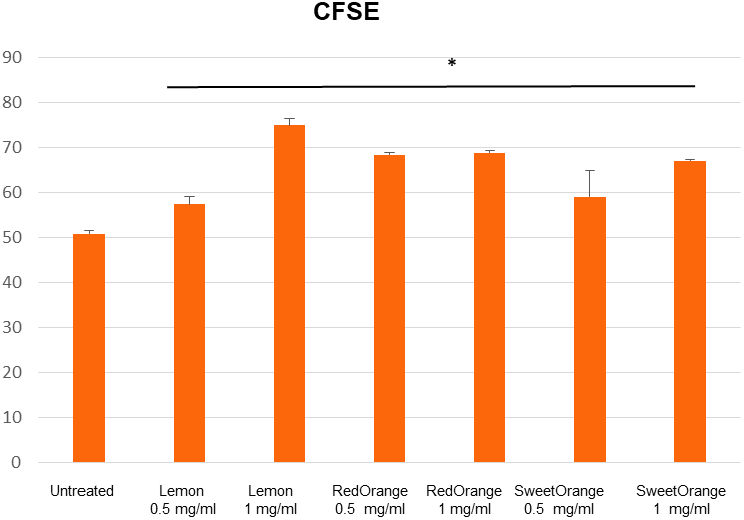
CFSE fluorescence intensity signals from A549 cells treated with lemon, red orange, and sweet orange IntegroPectin at different concentrations.

The dose-dependent, high CFSE signals from cells treated with lemon, red orange and sweet orange IntegroPectin bioconjugates indicates that the cells treated with IntegroPectin conjugates underwent fewer divisions. This outcome suggests that all three pectin-based IntegroPectin bioconjugates effectively limit cell proliferation, preventing the typical progression through multiple cycles of cell division.

### 2.3 Effects on oxidative stress (MitoSox and ROS) in A549 cell line

Dual assessment of ROS in both the cytoplasm and mitochondria is crucial, as it provides a comprehensive understanding of the cellular oxidative environment. A549 cells were thus stimulated for 3 h with different IntegroPectin conjugates in solution at 0.5 and 1.0 mg/mL concentration and MitoSox and ROS assessed by cytofluorimetric analysis.

Plots in Fig.6 and Fig.7 show that at all tested concentrations, the IntegroPectin bioconjugates significantly reduced oxidative stress when compared to untreated cells, as clearly demonstrated by the marked decrease in both mitochondrial (A) and cytoplasmic (B) ROS levels.

**Figure 6.**
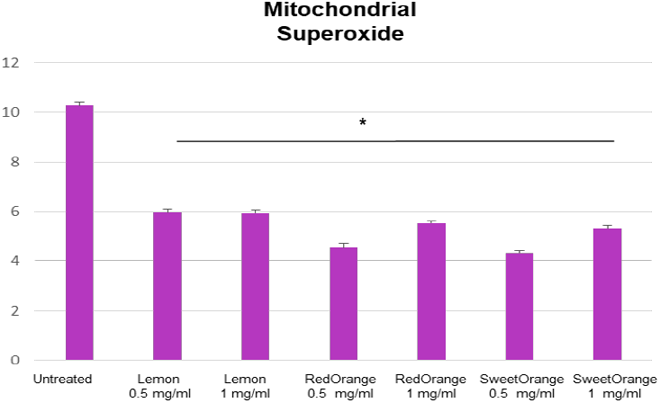
Mitochondrial oxidative stress in A549 cells treated with lemon, red orange, sweet orange IntegroPectin at different concentrations compared to untreated cells.

**Figure 7.**
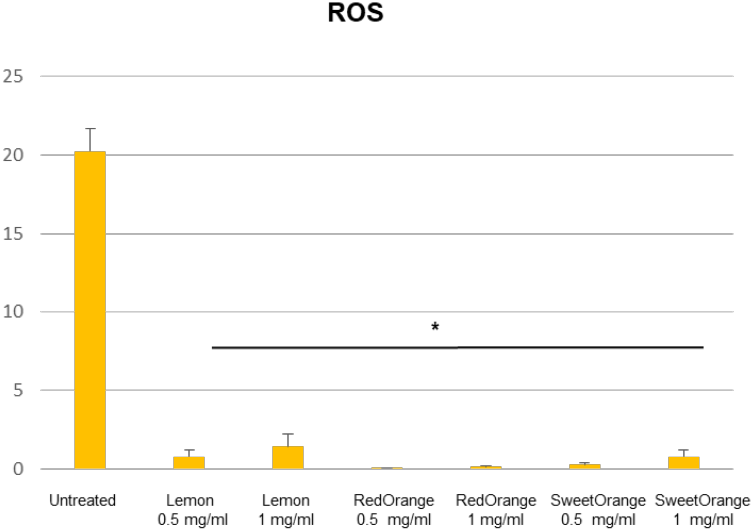
Cytoplasmic ROS levels in A549 cells treated with lemon, red orange, sweet orange IntegroPectin at different concentrations compared to untreated cells.

The reduction in oxidative stress suggests that IntegroPectin phytocomplexes help alleviate cellular oxidative damage restoring a more balanced cellular environment

### 2.4 Effects on miR-146 expression in A549 cell line

The expression of miR-146a was evaluated in A549 cell line treated with IntegroPectin extracted from lemon, red orange, and sweet orange. A concentration of 1 mg/mL was selected for these treatments, based on previous data demonstrating a significant pro-apoptotic effect at this dose.

Real-time quantitative polymerase chain reaction (RT-qPCR) analysis revealed that all three types of IntegroPectin induced an upregulation of miR-146a expression compared to untreated controls (Fig.8), with a substantially more significant effect of orange and red orange IntegroPectin bioconjugates.

**Figure 8.**
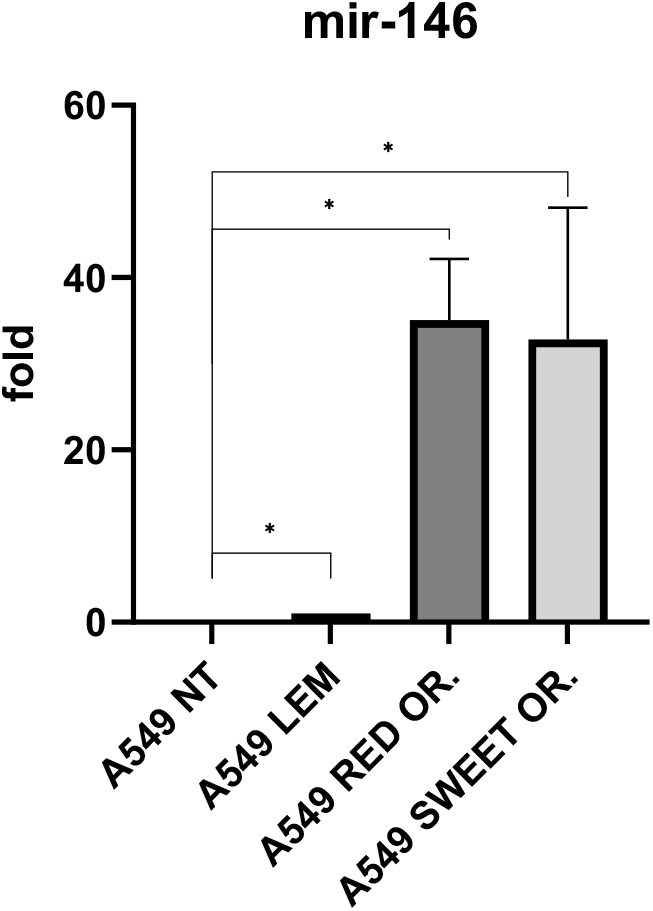
Upregulation of miR-146a expression driven by lemon, red orange and sweet orange IntegroPectin bioconjugates in lung adenocarcinoma A549 cells compared to untreated control.

This increase suggests a potential therapeutic effect of the flavonoid-pectin bioconjugates under investigation. miR-146, particularly miR-146a, is a key regulator of inflammatory responses and cellular stress pathways. In cancer models, it has been shown to modulate critical processes such as apoptosis, proliferation, and immune evasion, often acting as a tumor suppressor by targeting components of the NF-κB signaling pathway, such as IRAK1 and TRAF6.^[18,19]^ Its upregulation is associated with reduced tumor growth and metastatic potential in various malignancies, including breast and lung cancers.^[20,21]^

The observed increase in miR-146 expression in A549 cells upon *Citrus* IntegroPectin treatment supports the hypothesis that these natural compounds may exert anti-tumoral effects via modulation of microRNA networks involved in cancer progression.

### 2.5 Effects on IL-8 expression in A549 cell line

Interleukin-8 (IL-8 or CXCL8) is a pro-inflammatory chemokine promoting tumor growth, angiogenesis, and metastasis.^[19]^ Hence, we evaluated the expression of IL-8 mRNA in A549 cell line treated with IntegroPectin extracted from lemon, red orange, and sweet orange.

Treatment with 1 mg/mL of lemon, red orange, or sweet orange IntegroPectin resulted in a decrease of IL8 expression in A549 cells (Fig.9).

**Figure 9.**
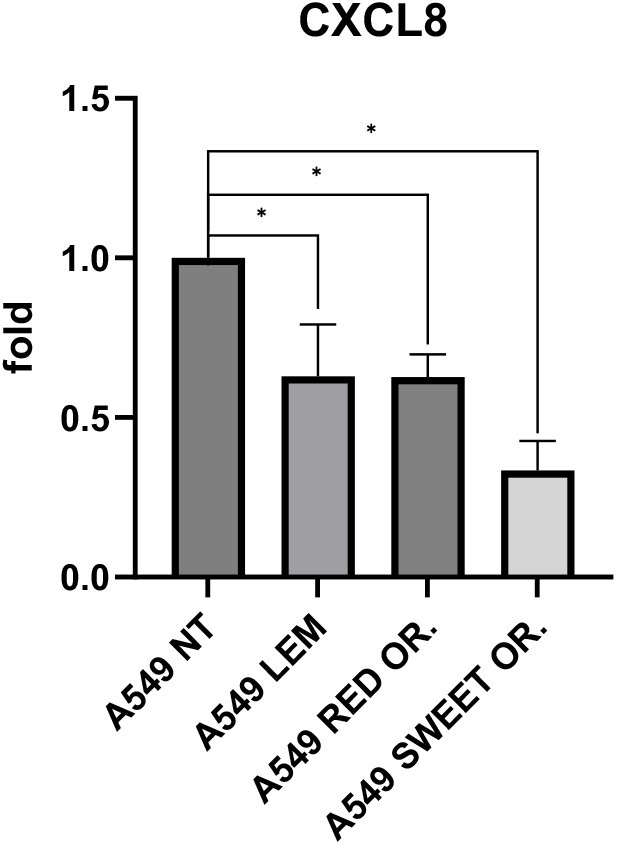
Downregulation of IL-8 mRNA driven by lemon, red orange and sweet orange IntegroPectin bioconjugates in lung adenocarcinoma A549 cells compared to untreated control.

The decrease in IL-8 mRNA levels, with the concomitant increase in miR-146a expression, supports the hypothesis that IntegroPectin may exert an anti-tumoral effects, through the induction of miR-146a, which is known to target upstream regulators of IL8 transcription. Besides driving apoptosis and inhibiting cancer cell growth and migration, in lung cancer miR-146a delays EMT and attenuates IL 1β induced IL-8 secretion.^[22]^

The inverse relationship between miR-146a and IL-8 expression suggests that compounds capable of upregulating miR-146a may also suppress IL-8 expression and its tumor-promoting effects.^[23]^

In brief, these results show that treatment with sweet orange, red orange and lemon IntegroPectin bioconjugates in solution induces an upregulation of miR-146 and a downregulation of IL-8 expression, suggesting that *Citrus* IntegroPectins bioconjugates may contribute to anti-tumorigenic activity through miRNA-mediated modulation of cancer-related signaling pathways in the tumor microenvironment through an epigenetic mechanism.

## 3. Discussion

Three main results of this *in vitro* investigation using adenocarcinoma A549 cell line provide new insights into the lung anticancer activity of *Citrus* IntegroPectin bioconjugates sourced via cavitation of industrial lemon, red orange, and sweet orange industrial biowaste originating from organically grown fruits.

First, driving apoptosis, these pectins impact the altered apoptosis and uncontrolled cellular growth characteristics of all cancerous diseases, leading to uncontrolled cancer cells growth, allowing the accumulation of mutations which are the cause of angiogenesis, invasiveness and cell proliferation. Remarkably, therapeutic advancements in cancer treatment in the past 25 years saw the emergence of small-molecule targeted therapy and immunotherapy treatments, both of which exert their antitumor effects by triggering apoptosis.^[24]^

In detail, treatment of A549 cells with IntegroPectin conjugates, particularly with lemon IntegroPectin at 1.0 mg/mL and red orange IntegroPectin at both 0.5 and 1.0 mg/mL concentration, resulted in a significant increase in apoptotic cell death. The apoptotic effect appears to be dose-dependent, with higher concentrations leading to stronger apoptosis, which may be indicative of a potent anticancer effect of these compounds.

This apoptotic response, furthermore, is accompanied by a cell cycle arrest at the G0/G1 phase. The G0/G1 phase is a critical checkpoint in the cell cycle, where cells either remain in a quiescent state (G0) or prepare to enter the S phase for DNA replication (G1). The arrest at this stage effectively blocks the progression of tumor cells into the DNA synthesis (S phase) and mitosis (G2/M phase) By halting progression through the cell cycle at an early stage, IntegroPectin phytocomplexes essentially prevent cancer cells from proliferating and dividing, thus reducing overall tumor growth. This suggests that IntegroPectin conjugates may have a cytostatic effect, particularly in inhibiting cell cycle progression at a crucial phase limiting the ability of the A549 cells to proliferate, thus preventing the typical progression through multiple cycles of division.

Notably, this behavior of hesperidin-rich red orange and sweet orange IntegroPectin bioconjugates is similar to the G0/G1 arrest of A549 cells caused by hesperidin dissolved in DMSO, driven by induced apoptosis through the mitochondrial apoptotic pathway.^[25]^

Second, all (lemon, red orange and sweet orange) *Citrus* IntegroPectin biooconjugates investigated substantially reduce oxidative stress levels virtually eliminating cytoplasmic ROS levels and substantially reducing mitochondrial superoxide. At the cellular level, ROS originate from multiple sources. Mitochondria are a major site of ROS production through the electron transport chain, generating superoxide and hydrogen peroxide locally. This compartmentalized production can lead to specific damage to mitochondrial DNA, proteins, and lipids, which would be missed if only cytoplasmic ROS were assessed.^[26]^ Importantly, mitochondrial ROS levels are widely used as markers of mitochondrial health and dysfunction, which are implicated in numerous diseases, including neurodegeneration, diabetes and cancer.^[27]^

The large reduction in both cytoplasmic and mitochondrial ROS is in agreement with the pronounced antioxidant activity of *Citrus* IntegroPectin bioconjugates,^[9]^ in its turn is due to the substantial amounts of flavonoids chemically bound to the structure of the LM pectin.^[28]^

Basic research and clinical trials in lung cancer therapy have demonstrated that increasing ROS has anticancer potential, whereas scavenging ROS even promotes metastasis.^[29]^ However, the powerful antioxidant and radical scavenging activity of the IntegroPectin bioconjugates ensures elimination of oxidative stress (the beneficial effect of antioxidants) without being detrimental to the ability of these pectins to induce apoptosis.

Indeed, the third main finding of this work is that at the concentration of 1.0 mg/mL we observed significant upregulation of miR-146 and downregulation of IL-8 cytokine, one of its downstream targets. Enhanced miR-146 expression in A549 lung cancer cells, indeed, increases apoptosis and autophagy by targeting the TNF receptor-associated factor-6 (TRAF6) gene involved in apoptosis and affecting other genes such as BCL-2, IL-6, and TNFα through the NF-kB signaling pathway.^[30]^

Downregulation of pro-inflammatory Interleukin-8, in its turn, is highly beneficial as IL-8 is a growth factor for human lung cancer cells^[31]^ that inhibits apoptosis and promotes EMT, cell migration, invasion, angiogenesis and leads to metastasis in numerous cancers, including lung cancer.^[32]^ Indeed, IL-8 is now a pharmacological target for the optimization of therapies, including immunotherapeutic approaches.^[33]^ Reduction of IL-8, in brief, removes key anti-apoptotic signals, sensitizing cells to apoptotic stimuli.

Citrus flavonoids including hesperidin,^[25]^ naringenin,^[34]^ eriocitrin,^[35]^ and kaempferol^[36]^ are widely investigated for the treatment of lung cancer. Research on the anticancer properties of depolymerized and de-esterified citrus pectin, in its turn, has clearly established that modified citrus pectin is both a galectin-3^[37]^ and galectin-8 inhibitor.^[38]^ Associated with several cancer types, galectin-1, −3, −4, −7, −8 and −9 in lung cancer are associated with tumor invasion, migration, metastasis and progression.^[39]^ Galectin-3 promotes cancer progression by attaching cancer cells to each other and helps in migration to other parts of the body.

Both free carboxylate and neutral sugars galactose and arabinose in pectin RG-I and RG-II regions interact with galectin-3 binding protein expressed on cancer cells and thus prevent the aggregation of cancer cells exerting thereby antiproliferative activity.^[40]^

Both low degree of esterification and higher abundance of galactan-rich RG-I side chains in *Citrus* IntegroPectin are markedly important structural features for enhanced bioactivity.^[9]^

Given the high biocompatibility and health beneficial properties of citrus pectin^[41]^ and citrus flavonoids,^[42]^ as well as the frequency and high mortality of lung cancer, these results further support urgent investigation of IntegroPectin bioconjugates in preclinical and clinical cancer treatment trials in the treatment of lung cancer.

Biomedical use of citrus flavonoids so far has been hindered by ultralow solubility and poor bioavailability of most citrus flavonoids.^[43]^ Now, hesperidin bound to highly biocompatible, readily soluble and health-beneficial citrus pectin at an unprecedented high load exceeding 62 mg/g such as in the case of (dialyzed) red orange IntegroPectin unveiled in this study, could be readily administered orally or injected along with other bioactive citrus flavonoids such as eriocitrin and kaempferol concomitantly present in these highly de-esterified pectin bioconjugates.^[9]^

## 4. Conclusions

In summary, following the discovery that *Citrus* IntegroPectin bioconjugates obtained through acoustic cavitation in water only of lemon, red orange, and sweet orange industrial processing waste show substantial anticancer action *in vitro* against human non-small cell lung cancer cells of line A549, we investigated the mechanism of action using IntegroPectin bioconjugates further purified from residual sugars via simple dialysis.

New results show that the anticancer properties of these highly bioactive bioconjugates originate by their ability to induce apoptosis, decrease short-term proliferation response, reduce ROS and alleviate mitochondrial oxidative stress. The same IntegroPectin bioconjugates induce also upregulation of miR146 expression and downregulation of Interleukin-8 expression.

These results provide further evidence that these highly biocompatible bioconjugates show remarkable pleiotropic activity in their cancer mechanism by targeting different molecular pathways. Their combined biochemical, cellular, and epigenetic effects underscore their potential for development as new anti-cancer therapy. Further supporting the aforementioned need for preclinical and clinical studies, modified citrus pectin has already used in various clinical applications, with recent results (2020-early 2025) supported by multiple *in vitro* and *in vivo* studies, as well as early-phase clinical trials confirming that said pectin has enhanced bioavailability and significant anticancer activity.^[44]^

## 5 Materials and methods

### 5.1 IntegroPectin isolation

Three *Citrus* fruit IntegroPectin bioconjugates were obtained via acoustic cavitation of fresh citrus processing waste. In detail, lemon, sweet orange, and red orange CPW obtained from industrial citrus fruit squeezing in Sicily was kindly provided by OPAC Campisi (Siracusa, Italy). The CPW was packed in cardboard boxes and stored in a cold chamber at 4 °C during the transportation. All raw CPW samples were stored in a freezer at –20 °C and brought to room temperature prior to the AC-assisted extraction.

An aliquot (300 g) of CPW at room temperature was added with 3 L of ultrapure water obtained using a Barnstead Smart2Pure Water Purification System (Thermo Fisher Scientific, Waltham, MA, USA) and homogenized with a domestic electric blender by grinding twice for 30 s at high speed each time. The resulting mixture was extracted using the UIP2000hdT (20 kHz, 2000 W) industrial sonicator (Hielscher Ultrasonics, Teltow, Germany) equipped with a hydraulic pump operating at 1.43 L/min. The extraction process was carried out in continuous flow-mode for 30 min at 50% of amplitude, in pulse condition (50 s on 50 s off), setting the maximum work temperature at 50 °C.

The power supplied to the digital probe-type sonicator was set at 800 W. After extraction was complete, the mixture was filtered through a cotton cloth in order to separate the insoluble fraction from the aqueous phase. The aqueous phase containing the IntegroPectin in solution was further filtered through a Büchner funnel by passing the mixture through a filter paper (Whatman, grade 589/3, retention < 2 μm) placed in the funnel.

Each type of IntegroPectin (lemon, sweet orange, and red orange) was dispersed in 5 mL of distilled water until a clear solution was obtained. The sample was then transferred into dialysis bag (12.5 kDa cut-off, Millipore) immersed in 50 mL of distilled water. The filled bag was kept under magnetic stirring for 24 h. After the first hour, the entire acceptor fluid was replaced with fresh distilled water to accelerate diffusion of residual sugar molecules. At the end of the dialysis process, each IntegroPectin bioconjugate was isolated by freeze-drying for 48 each IntegroPectin in solution retrieved from the dialysis bag using a FreeZone 4.5 Liter Benchtop Freeze Dry System (Labconco, Kansas City, MO, USA). The resulting dry materials were collected and stored at 4 °C until further characterization. Flavonoids quantitative evaluation

### 5.2 FTIR analyses

The FTIR (Fourier Transform Infrared) spectroscopy measurements were carried out using an Alpha spectrometer (Bruker, Billerica, MA, USA). A sample (2 mg) of each IntegroPectin was mixed with ultrapure KBr (100 mg, FTIR grade, ≥99% trace metals basis, Sigma-Aldrich, St. Louis, MI, USA). The mixed powder was ground using a pestle in an agate mortar to form a uniform mixture. A Specac Mini-Pellet laboratory hydraulic press was used for the preparation of high-quality 7 mm KBr pellets for transmission FTIR applying a 10 tonne weight for 3 min. Spectra were collected over 400–4000 cm−^1^ at a resolution of 2 cm−^1^, with 128 scans per spectrum to improve the signal-to-noise ratio. A background spectrum recorded under identical conditions was automatically subtracted to minimize contributions from atmospheric components.

### 5.3 Flavonoid quantification

A 10 mg sample of each IntegroPectin was dispersed in 1 mL of DMSO using a sonication bath. The resulting dispersion was then filtered through a 0.22 µm nylon membrane filter and analyzed by high-performance liquid chromatography coupled with a diode array detector (HPLC-DAD). In detail, chromatographic separation was performed on a 1260 Infinity HPLC system (Agilent Technologies, Santa Clara, CA, USA) equipped with a binary pump and a diode array detector. A Synergy Hydro-RP C18 column (150 mm × 4.6 mm; 80 Å pore size; 4 μm particle size) purchased from Phenomenex (Torrance, CA, USA) served as the stationary phase.

The following conditions were employed: mobile phase A—water with 0.1% trifluoroacetic acid (TFA); mobile phase B—acetonitrile with 0.1% TFA. The elution gradient was as follows: 0– 2 min, 90:10 (A/B); 2–27 min, 50:50 (A/B); 27–29 min, 50:50 (A/B); 29–31 min, 90:10 (A/B). The flow rate was set at 1.5 mL/min, the column temperature was maintained at 35 °C, and the detection wavelength was set at 285 nm. Quantification of flavonoids was achieved using calibration curves constructed from five standard solutions for each analyte. All analyses were conducted in triplicate, and flavonoid content was expressed as mg/g of each IntegroPectin, reported as mean ± standard deviation (SD, *n*=3). Results were expressed as mg of each flavonoid per g of IntegroPectin.

### 5.4 IntegroPectin solutions in PBS

A sample of each IntegroPectin was dispersed in PBS (phosphate-buffered saline, pH 7.4, purchased from Gibco Invitrogen, New York USA) at a concentration of 20 mg/mL. The resulting mixture was sonicated for 3 min to achieve a homogeneous solution. For each IntegroPectin, the resulting solution was stored at 4 °C prior to the biological tests.

### 5.5 Cell Culture and Treatment

Non-small cell lung cancer (NSCLC) adenocarcinoma cell line (A549 CCL-185™), were cultured in a humidified ambient and air with 5 % CO_2_ at 37 °C. Cells were grown as adherent monolayers in RPMI 1640 medium supplemented with heat deactivated (56 °C, 40 min) 10 % FBS, 50 U/mL penicillin, 50 mg/mL streptomycin, 1 % non-essential amino acids and 2 mM L-glutamine (all from Euroclone, Pero, Italy). Cells were cultured to confluence and stimulated, for the indicated time, with each of the three IntegroPectin samples dissolved in PBS at 0.5 and 1.0 mg/mL concentration and then harvested for further investigation. At least three replicates were performed for each experiment.

### 5.6 Cell Cycle and Apoptosis Analysis

A549 were seeded in 12-well plate and stimulated as already described for 24 h. After treatment, cells were harvested, washed with ice-cold PBS, and resuspended at a concentration of 1 × 10^6^ cells/mL in a hypotonic fluorochrome solution containing 0.1% sodium citrate, 0.03% Nonidet P-40, and 50 μg/mL propidium iodide. The cells were incubated for 30 min at room temperature in the dark Cell acquisition was performed using the CytoFLEX system (Beckman Coulter, Lakeview Pkwy S Drive, Indianapolis, USA). For each sample, 10,000 events were acquired. Data were analyzed using CytExpert software. Based on DNA content, cells were classified as follows: P1 (apoptotic cells), P2 (sub-G1 phase), P3 (G0/G1 phase), P4 (S phase), and P5 (G2/M phase). Results were expressed as the percentage of cells in each population

### 5.7 CFSE Labeling Assay

Cell proliferation was assessed using the carboxyfluorescein succinimidyl ester (CFSE) labeling assay). In detail, A549 cells were labeled with CFSE (Molecular Probes, Eugene, OR, USA) at a final concentration of 5 μM and incubated at 37 °C for 10 min. The staining reaction was stopped by adding an equal volume of fetal bovine serum and mixing thoroughly. Cells were then centrifuged, collected, and washed twice with cold PBS to remove any unbound CFSE. After washing, cells were resuspended in complete medium, seeded into culture plates, and stimulated for 24 h with different *Citrus* IntegroPectin solutions, as already described. After 72 h, the A549 cells were harvested, washed with PBS and CFSE fluorescence intensity was measured by flow cytometry using the CytoFLEX system (Beckman Coulter, Lakeview Pkwy S Drive, Indianapolis, USA). For each sample, 10,000 events were acquired and analyzed using CytExpert software. Results were expressed as the percentage of CFSE-positive cells. A reduction in CFSE intensity indicated cell proliferation, because when a cell divides, the CFSE is distributed equally between the two daughter cells, resulting in a halving of the fluorescence intensity in each daughter cell compared to the parent cell.

### 5.8 Mitochondrial Superoxide Production (MitoSOX)

Mitochondrial superoxide production was assessed by flow cytometry using the MitoSOX Red mitochondrial superoxide indicator (Molecular Probes, Waltham, MA, USA). The A549 cells were seeded in 12-well plate and stimulated as already described for 3 h. Then the cells were harvested, washed with PBS 1% FBS and stained with 3 μM MitoSOX Red for 15 min at 37 °C. Subsequently, the cells were washed twice with PBS 1% FBS and analyzed by flow cytometry using the CytoFLEX system (Beckman Coulter, Indianapolis, USA). For each sample, 10,000 events were acquired. Data were analyzed using CytExpert software, and results were expressed as the percentage of MitoSOX-positive cells.

### 5.9 Intracellular Reactive Oxygen Species (ROS)

Intracellular ROS levels were measured by flow cytometry using the ROS-sensitive probe 6-carboxy-2′,7′-dichlorodihydrofluorescein diacetate (H2DCFDA, C-2938 Invitrogen by Thermo Fisher Scientific). A549 cells were seeded in 12-well plates and stimulated as previously described for 3 hours. Subsequently, the cells were harvested, washed with PBS, and stained with 0.5 μM H2DCFDA for 30 minutes at room temperature in the dark. After incubation, the cells were washed again with PBS and analyzed by flow cytometry using the CytoFLEX system (Beckman Coulter, Indianapolis, USA). For each sample, 10,000 events were acquired. Data analysis was performed using CytExpert software, and results were expressed as the percentage of ROS-positive cells.

### 5.10 RNA Extraction

The total RNA was extracted by A549 cell line, treated or not with IntegroPectin samples dissolved in PBS at 1 mg/mL, using RNAspin mini kit (GE Healthcare Science, Uppsala, Sweden), according to the manufacturer’s instruction. The RNA concentration was assessed using park multimode microplate reader (Tecan, Switzerland). For this study, only RNA with a ratio of A260/280 from 1.9 to 2 was employed.

### 5.11 TaqMan RT-qPCR for miR-146a-5p and IL8

The cDNA was synthesized from total RNA, using iScript cDNA synthesis kit (Biorad, Hercules, CA, USA), according to the manufacturer’s protocol. IL8 gene expression was evaluated with specific FAM-labelled probe and primers, with IL8 TaqMan gene expression assay (Hs99999034_m1 Applied Biosystems, Foster City, CA, USA), by real-time quantitative PCR (RT-qPCR) using QuantStudio Real-Time PCR system (Applied Biosystems, Foster City, CA, USA). The gene expression was normalized to GAPDH by GAPDH Taq-Man gene expression assay (Hs03929097_g1, Applied Biosystems, Foster City, CA, USA) as a housekeeping gene.

For amplification, the reaction mixtures were incubated at 95 °C for 15 min, followed by 40 amplification cycles of 94 °C for 15 s, 55 °C for 30 s, and 70 °C for 30 s. The reverse transcription of miR-146a-5p, 0.5 µg of total RNA was performed with specific primers using TaqMan miRNA RT kit (Applied Biosystems, Foster City, CA, USA), according to the manufacturer’s protocol. The miR-146a-5p expression was evaluated using specific TaqMan microRNA (Assay ID 000468, Applied Biosystems), with RTqPCR by QuantStudio Real-Time PCR system (Applied Biosystems). The miR-146a-5p expression in A549 cells was normalized with RNU6 using the TaqMan microRNA assay (Assay ID 001973, Applied Biosystems). Triplicate samples and inter-assay controls were used. The 2-ΔCT method was employed for the normalization of RT-qPCR data.

### 5.12 Statistical Analysis

Analysis of variance (ANOVA) was used to assess differences between group means. Associations between categorical variables were evaluated using Fisher’s exact test. A value of *p*< 0.05 was considered statistically significant.

## Acknowledgements

We thank OPAC Campisi (Siracusa, Italy) for the generous gift of industrial processing waste of organically grown citrus fruits from which the IntegroPectin bioproducts were sourced. Work of G.L.P. was supported by MICS (Made in Italy - Circular and Sustainable) Extended Partnership and received funding from the European Union NextGenerationEU (PNRR - Mission 4 Component 2, Investment 1.3 - D.D.1551.11-10-2022, PE00000004). Work of G.A. was supported by the SAMOTHRACE (Sicilian Micro and Nano Technology Research and Innovation Center) Innovation Ecosystem using funding from European Union NextGenerationEU (PNRR – Mission 4 Component 2, Investment 1.5 (ECS00000022)). R.C. and M.P. thank Ministero dell’Università e della Ricerca for funding, under Progetto “FutuRaw. Le materie prime del futuro da fonti non-critiche, residuali e rinnovabili”, Fondo Ordinario Enti di Ricerca, 2022, (CUP B53C23008390005).

## Data availability statement

The data that support the findings of this study are available from the corresponding authors upon reasonable request.

---

Tested *in vitro* on adenocarcinoma A549 cell line, IntegroPectin bioconjugates derived from the industrial processing waste of different organically grown *Citrus fruits*, exert a pleiotropic activity in their anticancer mechanism by targeting different molecular pathways. This study shows that these novel bioconjugates induce apoptosis, decrease short-term proliferation response, reduce reactive oxygen species (ROS), alleviate mitochondrial oxidative stress, upregulate miR-146 expression and downregulate expression of Interleukin-8. Given the health beneficial properties of both citrus pectin and citrus flavonoids, and the frequency of lung cancer, said findings support urgent investigation of these flavonoid-pectin conjugates in preclinical and clinical cancer treatment trials.

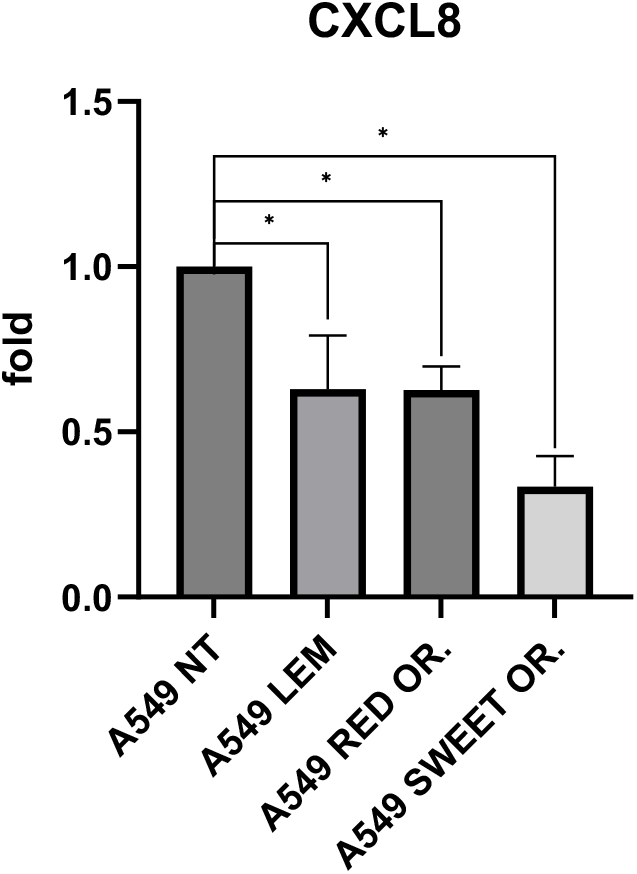

