## Supplemental Figures S1 and S2, Supplemental Table S1 for "*Citrus* IntegroPectin bioconjugates induce apoptosis, decrease proliferation, reduce reactive oxygen species, alleviate mitochondrial oxidative stress, upregulate miR-146 expression and downregulate expression of Interleukin-8 in lung cancer cells"

*^1^Istituto di Farmacologia Traslazionale, CNR, via U. La Malfa 153, 90146 Palermo, Italy; ^2^Istituto per lo Studio dei Materiali Nanostrutturati, CNR, via U. La Malfa 153, 90146 Palermo, Italy*

**Corresponding authors*

**Supplementary Information**

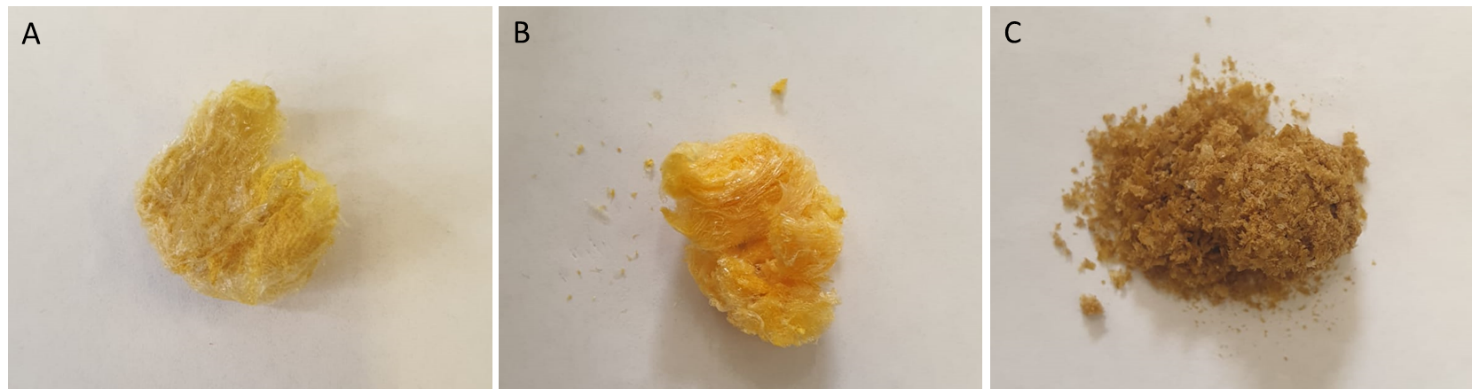

**Figure S1**. *Citrus* IntegroPectin samples freeze dried after 24 h dialysis:

A) lemon IntegroPectin, B) sweet orange IntegroPectin, C) Red orange IntegroPectin.

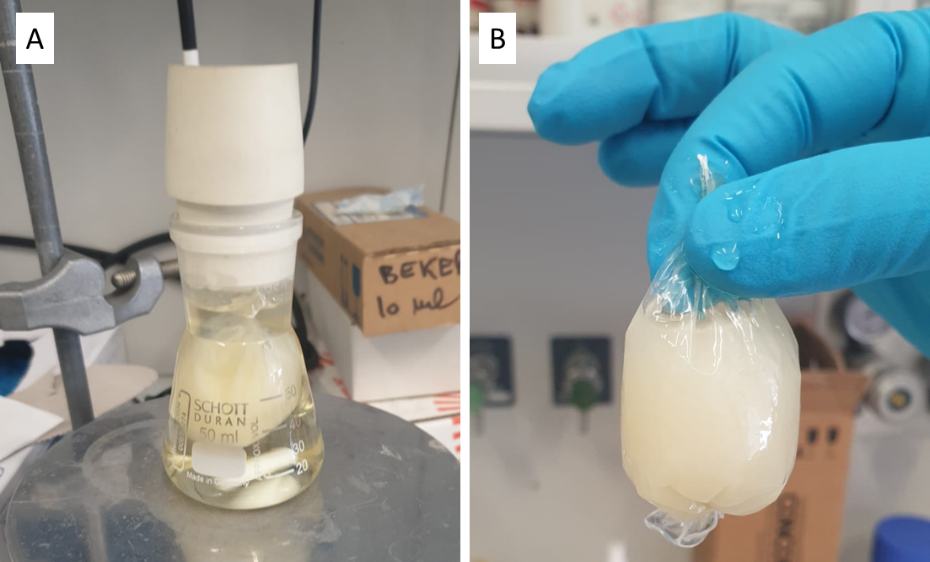

**Figure S2**. Purification of lemon IntegroPectin:

A) Dialysis setup showing lemon IntegroPectin in a dialysis membrane immersed in water.

B) Appearance of the purified lemon IntegroPectin after dialysis (24 h).

**Table S1**. Representative flavonoid (eriocitrin and hesperidin) content in dialyzed lemon, sweet and red orange IntegroPectin bioconjugates after 24 h dialysis.

| ***Citrus* IntegroPectin** | **Eriocitrin** (mg/g) | **Hesperidin** (mg/g) |
| --- | --- | --- |
| Lemon | 2.14 | 4.9 |
| Red orange | - | 62.31 |
| Sweet orange | 1.23 | 40.35 |
